# Domain-General and Domain-Specific Patterns of Activity Supporting Metacognition in Human Prefrontal Cortex

**DOI:** 10.1101/172445

**Authors:** Jorge Morales, Hakwan Lau, Stephen M. Fleming

## Abstract

Metacognition is the capacity to evaluate the success of one’s own cognitive processes in various domains, e.g. memory and perception. It remains controversial whether metacognition relies on a domain-general resource that is applied to different tasks, or whether self-evaluative processes are domain-specific. Here we directly investigated this issue by examining the neural substrates engaged when metacognitive judgments were made by human participants during perceptual and memory tasks matched for stimulus and performance characteristics. By comparing patterns of functional magnetic resonance imaging (fMRI) activity while subjects evaluated their performance, we revealed both domain-specific and domain-general metacognitive representations. Multi-voxel activity patterns in anterior prefrontal cortex predicted levels of confidence in a domain-specific fashion, whereas domain-general signals predicting confidence and accuracy were found in a widespread network in the frontal and posterior midline. The demonstration of domain-specific metacognitive representations suggests the presence of a content-rich mechanism available to introspection and cognitive control.

**Significance statement:** We use human neuroimaging to investigate processes supporting memory and perceptual metacognition. It remains controversial whether metacognition relies on a global resource that is applied to different tasks, or whether self-evaluative processes are specific to particular tasks. Using multivariate decoding methods, we provide evidence that perceptual- and memory-specific metacognitive representations cortex co-exist with generic confidence signals. Our findings reconcile previously conflicting results on the domain-specificity/generality of metacognition, and lay the groundwork for a mechanistic understanding of metacognitive judgments.

## Introduction

Metacognition is the capacity to evaluate the success of one’s cognitive processes in various domains, e.g. perception or memory (Flavell, 1979; Nelson and Narens, 1990; Metcalfe and Shimamura, 1994; Fleming et al., 2012a). Metacognitive ability can be assessed in the laboratory by quantifying the trial-by-trial correspondence between objective performance and subjective confidence (Galvin et al., 2003; Maniscalco and Lau, 2012; Overgaard and Sandberg, 2012; Fleming and Lau, 2014). Anatomical (Fleming et al., 2010; McCurdy et al., 2013; Maniscalco et al., 2017), functional (Fleck et al., 2006; Yokoyama et al., 2010; Fleming et al., 2012b; Baird et al., 2013; Hilgenstock et al., 2014) and neuropsychological (Shimamura and Squire, 1986; Schnyer et al., 2004; Fleming et al., 2014) evidence indicates specific neural substrates (especially in frontolateral, frontomedial, and parietal regions) contribute to metacognition across a range of task domains, including perception and memory. However, the neurocognitive architecture supporting metacognition remains controversial. Does metacognition rely on a common, domain-general resource that is recruited to evaluate performance on a variety of tasks? Or is metacognition supported by domain-specific components?

Current computational perspectives (Pouget et al., 2016; Fleming and Daw, 2017) suggest both domain-general and domain-specific representations may be important for guiding behavior. On the one hand, one needs to be able to compare confidence estimates in a “common currency” across a range of arbitrary decision scenarios (de Gardelle and Mamassian, 2014). One solution to this problem is to maintain a global resource with access to arbitrary sensorimotor mappings (Holroyd et al., 2005; Heekeren et al., 2006; Cole et al., 2013). Candidate neural substrates for a domain-general resource are the frontoparietal and cingulo-opercular networks, known to be involved in arbitrary control operations (Cole et al., 2013). In particular, the dorsomedial prefrontal cortex (encompassing the paracingulate cortex and pre-supplementary motor area) has been implicated in representing confidence, monitoring conflict, and detecting errors across a range of tasks (Gehring et al., 1993; Botvinick et al., 2004; Ridderinkhof et al., 2004; Fleming et al., 2012b). On the other hand, if the system only had access to generic confidence signals, appropriate switching between particular tasks or strategies on the basis of their expected success would be compromised. Functional imaging evidence implicates human anterior prefrontal cortex in tracking the reliability of specific alternative strategies during decision-making (Donoso et al., 2014), and such regions may also support domain-specific representations of confidence.

Current behavioral evidence of a shared resource for metacognition is ambiguous, in part due to the difficulty of distilling metacognitive processes from those supporting primary task performance (Galvin et al., 2003; Maniscalco and Lau, 2012; Fleming and Lau, 2014). Some studies have found that efficient metacognition in one task predicts good metacognition in another (McCurdy et al., 2013; Ais et al., 2016; Faivre et al., 2017; Ruby et al., 2017; Samaha and Postle, 2017), whereas others indicate the independence of metacognitive abilities (Kelemen et al., 2000; Baird et al., 2013; Vo et al., 2014). Recent studies employing bias-free measures of metacognition have identified differences in the neural correlates of memory and perceptual metacognition in both healthy subjects (Baird et al., 2013; McCurdy et al., 2013) and neuropsychological patients (Fleming et al., 2014). However, the study of behavioral individual differences provides only indirect evidence of the neural and computational architecture supporting metacognition.

Here, we directly investigate this ontology by examining neural substrates engaged when metacognitive judgments are made during perceptual and memory tasks matched for stimulus and performance characteristics. We employ a combination of univariate and multivariate analyses of functional magnetic resonance imaging (fMRI) data to identify domain-specific and domain-general neural substrates engaged during metacognitive judgments. We also distinguish activations engaged by a metacognitive judgment from neural activity which tracks confidence level. Together, our findings reveal the co-existence of generic and specific confidence representations, consistent with a computational hierarchy underpinning effective metacognition.

## Materials and Methods

### Participants

Thirty healthy subjects (ages 18-33, mean 24.97 years old; SD=4.44; 14 males) with normal or corrected-to-normal vision were monetarily compensated and gave written informed consent to participate in the study at the Center for Neural Science at New York University. The study protocols were approved by the local Institutional Review Board. The number of participants was determined a priori at n=30, which is in line with recent guidelines on neuroimaging sample sizes (Poldrack et al., 2017). Due to behavioral and in-scanner motion cut-off criteria, six subjects were excluded from further analysis (details below). We present the results of twenty-four subjects whose data were fully analyzed.

### Experimental and Task Design

The experiment had a 2×2×2 design: condition (confidence/follow) × task domain (perception/memory) × stimulus type (shapes/words). It consisted of six scanner runs, each with eight 9-trial mini-blocks (72 trials per run, 432 trials in total). Perceptual and memory two-alternative forced-choice (2-AFC) tasks were presented in separate, interleaved runs (three runs per task; order counterbalanced across subjects). In each run, there were four pairs of mini-blocks from the Confidence and Follow conditions. To avoid stimulus confounds, two different types of stimulus were used throughout the experiment. In each run, two pairs of Confidence/Follow mini-blocks used words and the remaining two pairs used abstract shapes (interleaved and order counterbalanced across runs).

In the perceptual task, subjects were asked to report the brighter of two stimuli on each trial. In the memory task, subjects began each mini-block by learning a set of nine consecutively-presented stimuli. A stimulus from this set was then presented on each subsequent trial (in randomized order) alongside a new stimulus. Subjects’ task was to identify the studied stimulus. In mini-blocks from the Confidence condition, subjects had to rate their confidence in their performance in each trial by selecting a number from a 1-to-4 scale. In mini-blocks from the Follow condition, subjects had to “follow the computer” in each trial by pressing the button corresponding to the highlighted number irrespective of their confidence. The highlighted number was yoked to their ratings in the previous Confidence mini-block (randomized presentation order) to ensure similar low-level visuomotor characteristics in both conditions for any given pair of mini-blocks.

Subjects were reminded at the beginning of each mini-block of the condition, task, and stimulus type that would follow. They used two fingers of their right hand to respond on an MRI-compatible button box: left stimulus (index) and right stimulus (middle). For confidence ratings, they used four fingers: 1 (index), 2 (middle), 3 (ring) and 4 (little). If subjects failed to provide either type of response within the allotted time (see Figure 1A for details), the trial was missed and an exclamation mark was displayed for the remainder of the trial. Failing to press the highlighted number counted as a missed trial.

**Figure 1.**
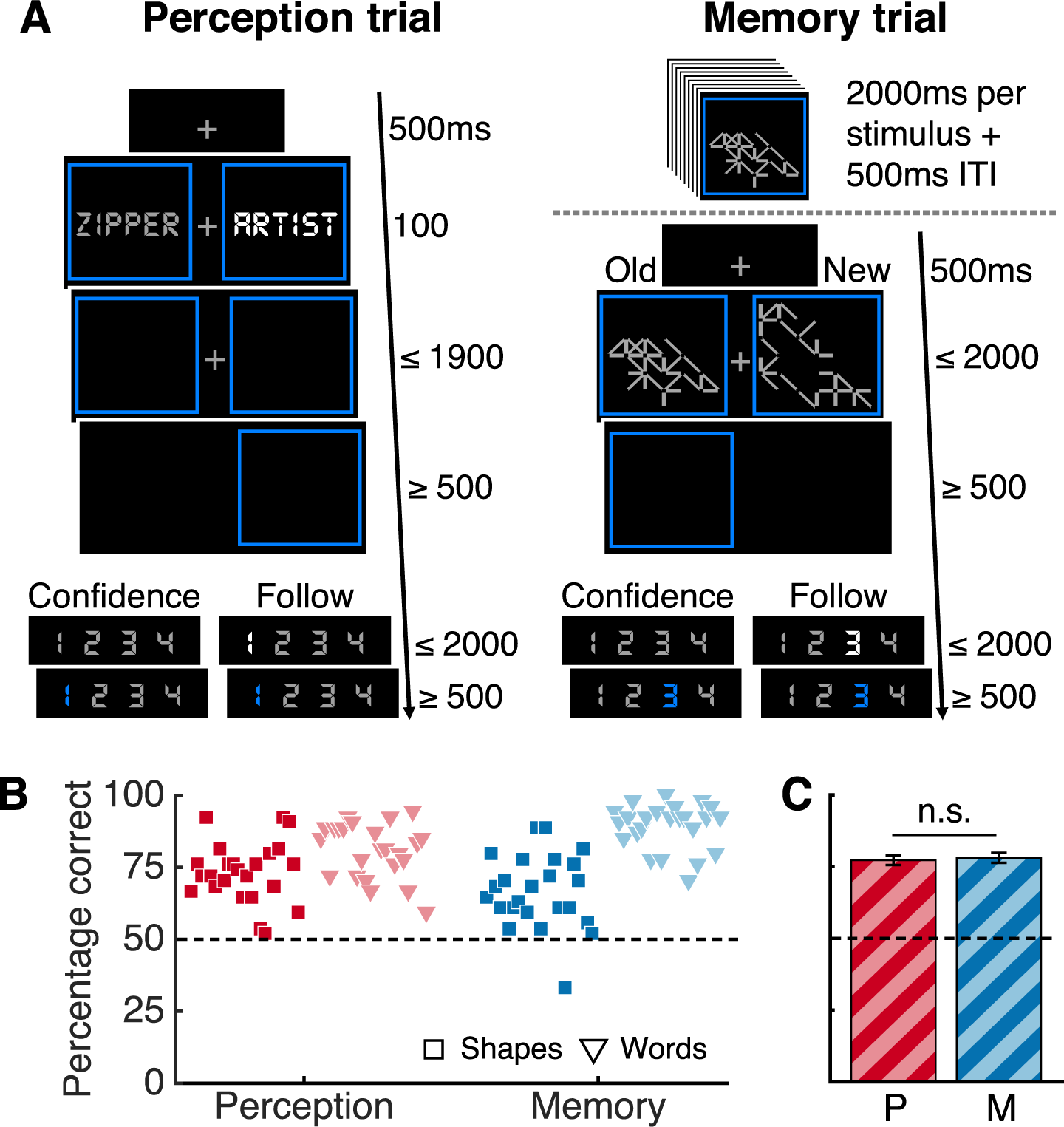
Task design & performance results. **(A)** Subjects performed two-alternative forced-choice discrimination tasks about perception and memory. In perception blocks, subjects selected the brighter of two stimuli. Memory blocks started with an encoding period and then subjects indicated in each trial which of two stimuli appeared during the encoding period. Abstract shapes and words were used as stimuli in both tasks. In Confidence blocks, subjects rated their confidence and in Follow blocks they pressed the highlighted number. **(B)** Percentage correct responses per block type in the Confidence condition. Each marker represents a subject. **(C)** Mean percentage correct responses by domain, averaged over subjects and stimulus types. Dotted lines indicate chance performance. Bars indicate standard error of the mean (s.e.m.). n.s.= not significant; P=perception; M=memory

Prior to entering the scanner, participants were familiarized with the tasks and the confidence rating scale. After computing independent brightness thresholding for words and abstract shapes, subjects practiced one of each mini-block type (i.e. 8 mini-blocks). Instructions emphasized that confidence ratings should reflect relative confidence and participants were encouraged to use all ratings. The whole experiment lasted ~1.5 hours.

### Stimuli

The experiment was programmed in MATLAB (MathWorks) and stimuli were presented using Psychtoolbox (Brainard, 1997). Abstract, 22- or 28-line shapes were randomly created by specifying an (invisible) grid of 6×6 squares that subtended 4 degrees of visual angle where lines could connect two vertices horizontally, vertically, or diagonally. The first line always stemmed from the central vertex of the invisible grid randomly connecting one of the surrounding eight vertices to ensure shape centrality within the grid. The remaining lines were drawn sequentially, ensuring all lines were connected. Orientation and originating vertices were selected randomly.

All words were nouns of 6 to 12 letters with 1 to 4 syllables obtained from the Medical Research Council Psycholinguistic Database (Wilson, 1988). In the perceptual task, words had high familiarity, concreteness, and imageability ratings (400-700). In the memory task, words had low ratings (100-400) to increase task difficulty. Each word and each shape was presented once throughout the experiment (across perceptual and memory blocks, including practice trials). All subjects were tested on the same words and shapes (counterbalanced across Confidence and Follow conditions across subjects). Words and rating scales were presented using DS-Digital font (40 points) to make their visual features similar to the abstract shapes.

To obtain stimulus sets of similar difficulty for shapes and words we ran a series of pilot studies where participants rated abstract shapes’ distinctiveness and then performed the memory task (15 mini-blocks per subject; 171 Amazon Mechanical Turk participants [73 for shapes; 98 for words] and 6 subjects in the laboratory who performed a complete version of the experiment). Based on these results, we expected a mean performance in the memory task of ~71% correct responses when 22- and 28-line distinctive shapes were used in the same block, and ~83% correct when long words (6-12 letters) with low concreteness, imageability and familiarity ratings (100-400) were used. To further increase difficulty, we created pairs of old and new words split between Confidence and Follow conditions (counterbalanced across subjects), blocked by similar semantic category (e.g. finance, argumentation, character traits, etc.), such that each new word within a block was freely associated with one old word (and when possible, vice versa) according to the University of South Florida free association normed database (Nelson et al., 2004).

In the perceptual task, the difference in brightness Δb) between the two stimuli was calibrated for each subject and independently for each stimulus type. The brightness of a randomly located reference stimulus was fixed (mid-gray). The brightness of the non-reference stimulus was titrated using a staircase procedure similar to previous experiments (Fleming et al., 2010; 2012b; 2014). During practice, we used a fixed large step size 2-down/1-up procedure until subjects reached 15 reversals or 90 trials. The step sizes followed recommended ratios to match the expected performance in memory blocks (García-Pérez, 1998). The experiment began with a Δb value determined by the average of the Δb values at each reversal, excluding the first one. Throughout the experiment, we kept a small step size staircase running to account for learning or tiredness.

A mid-gray fixation cross subtending 0.3 degrees of visual angle was presented between the two stimuli on a black background. The reference stimulus in the perceptual task and all stimuli in the memory task were mid-gray. All stimuli were surrounded by an isoluminant blue bounding box separated from the stimulus by a gap of at least 0.15 degrees of visual angle.

### Behavioral data analysis

Data analysis was performed in MATLAB and statistical analysis in RStudio (R Studio Team, 2015). We estimated metacognitive efficiency by computing log(meta-*d*′/*d*′). *d*′ is a signal detection theoretic measure of type 1 sensitivity, while meta-*d*′ is a measure of type 2 sensitivity (i.e. the degree to which a subject discriminates correct form incorrect responses) expressed in the same units as type 1 sensitivity (*d*′) (Maniscalco and Lau, 2012; Fleming and Lau, 2014). Meta-*d*’ indicates the *d*’ that would have been predicted to give rise to the observed confidence rating data assuming a signal detection theoretic ideal observer. Meta-*d*’=*d*’ indicates an optimal type II behavior for the observed type I behavior. Meta-*d*’ greater or less than *d*’ indicates metacognition that is better or worse, respectively, than expected given task performance, as may occur for instance if first-order decisions and confidence are supported by partly parallel processing streams (Fleming and Daw, 2017). We used hierarchical Bayesian estimation to incorporate subject-level uncertainty in group-level parameter estimates (Fleming, 2017). Certainty on this parameter was determined by computing the 95% high-density interval (HDI) from the posterior samples (Kruschke, 2010). For correlation and individual differences analyses we used single-subject Bayesian model fits. Two subjects were discarded for missing more than 10% of the trials (i.e. >1 standard deviation from the average missed trials, which was 5%). Missed trials were not analyzed.

### fMRI data acquisition

Brain images were acquired using a 3T Allegra scanner (Siemens). BOLD-sensitive functional images were acquired using a T2*-weighted gradient-echo echo-planar images (42 transverse slices, interleaved acquisition; TR, 2.34s; TE, 30ms; matrix size: 64×64; 3×3mm in-plane resolution; slice thickness: 3mm; flip angle: 90°; FOV: 126mm). The main experiment consisted of three runs of 210 volumes and three runs of 296 volumes for the perceptual and memory tasks, respectively. We collected a T1-weighted MPRAGE anatomical scan (1×1×1mm voxels; 176 slices) and local field maps for each subject.

### fMRI data preprocessing

Imaging analysis was carried out using SPM12 (Statistical Parametric Mapping; www.fil.ion.ucl.ac.uk/spm). The first five volumes of each run were discarded to allow for T1 stabilization. Functional images were realigned and unwarped using local field maps (Andersson et al., 2001) and then slice-time corrected (Sladky et al., 2011). Each participant’s structural image was segmented into gray matter, white matter, cerebral spinal fluid, bone, soft tissue, and air/background images using a nonlinear deformation field to map it onto template tissue probability maps (Ashburner and Friston, 2005). This mapping was applied to both structural and functional images to create normalized images to Montreal Neurological Institute (MNI) space.

Normalized images were spatially smoothed using a Gaussian kernel (8mm FWHM). We set a within-run 1mm rotation and 4mm affine motion cut-off criterion, which led to the exclusion of 4 subjects, leaving a total of 24 subjects whose functional and behavioral data were fully analyzed.

### Univariate analysis

All our general linear models (GLMs) focus on the ‘rating period’ of each trial by specifying boxcar regressors beginning at the subjects’ type I response and ending at their type II response (i.e. either confidence rating or number press). Motion correction parameters were entered as covariates of no interest along with a constant term per run. Regressors were convolved with a canonical hemodynamic response function (HRF). Low-frequency drifts were excluded with a 1/128Hz high-pass filter. Missed trials were not modeled. For judgment-related (JR) analyses, we created a GLM with two regressors of interest per run to estimate BOLD response amplitudes in each voxel during the ‘rating period’ in each trial of the Confidence and Follow blocks. For the confidence level-related (CLR) parametric modulation analysis, a GLM was used to estimate BOLD responses in Confidence blocks. There were two regressors of interest in each run, one modeling the confidence ‘rating period’ and another that encoded a parametric modulation by the four available confidence ratings (1-4).

#### Statistical inference

For the JR-analysis, single-subject contrast images of the Confidence and Follow regressors were entered into a second-level random effects analysis using one-sample *t*-tests against zero to assess group-level significance. For the CLR-parametric modulation analysis, single-subject contrast images of the parametric modulator were entered into a similar second-level random effects analysis. For conjunction analyses of activations common to both domains, second-level maps thresholded at *P*<0.001 (uncorrected) were intersected to reveal regions of shared statistically significant JR- and CLR-activity. Activations were visualized using MRIcro (http://www.mccauslandcenter.sc.edu/crnl/mricro). All second-level unthresholded statistical images were uploaded to Neurovault (Gorgolewski et al., 2015) (https://neurovault.org/collections/3232/).

#### ROI analysis

To define regions of interest (ROIs), 12mm spheres were centered at MNI coordinates identified from previous literature (Figure 3C). ROIs in left rostrolateral prefrontal cortex (L rlPFC) [−33, 44, 28], right rlPFC (R rlPFC) [27, 53, 25] and dorsal anterior cingulate cortex/pre-supplementary motor area (dACC/pre-SMA) [0, 17, 46] were created based on (Fleming et al., 2012b). The mask for precuneus (PCUN) [0, −64, 24] was based on (McCurdy et al., 2013). The MNI x-coordinates for the dACC/pre-SMA and PCUN masks were set to 0 to ensure bilaterality. Beta values were extracted from subjects’ contrast images for the JR- and CLR-univariate analyses, respectively.

### Multi-voxel pattern analysis

Multi-voxel pattern analysis (MVPA) was carried out in MATLAB using The Decoding Toolbox (Hebart et al., 2015). We classified run-wise beta images from GLMs modeling JR- and CLR-activity patterns in ROI and whole-brain searchlight analyses. ROI MVPAs were performed on normalized, smoothed images using the ROI spheres as masks. Previous work has shown that these preprocessing steps have minimal impact on support vector machine (SVM) classification accuracy, while allowing meaningful comparison across subject-specific differences in anatomy, as in standard fMRI analyses (Kamitani and Sawahata, 2010; Op de Beeck, 2010). A single accuracy value per subject, per condition and per ROI was extracted and used for group analysis and statistical testing. As a control, we added a 6mm-radius sphere centered at the ventricles [0 2 15].

Whole-brain searchlight analyses used 12mm-radius spheres centered around a given voxel, for all voxels, on spatially realigned and slice-time corrected images from each subject to create whole-brain accuracy maps. For group-level analyses, these individual searchlight maps were spatially normalized and smoothed using a Gaussian kernel (8mm FWHM) and entered into one-sample *t*-tests against chance accuracy (Hebart et al., 2014; 2015). Whole-brain cluster inference was carried out in the same manner as in univariate analysis. Visualizations were made with Surf Ice (https://www.nitrc.org/projects/surfice/).

Prior to decoding, for JR-activity pattern classification, we modeled two regressors of interest per run focused on the ‘rating periods’ in the Confidence and Follow conditions. For classification of CLR-activity patterns, we collapsed ratings 1 and 2 into a low-confidence regressor and ratings 3 and 4 into a high-confidence regressor to allow binary classification. The remaining parameters of no interest were specified as in the univariate case. For the CLR-searchlight analysis, we used an exclusive mask of activity patterns associated with usage of the confidence scale obtained from the successful cross-classification of button presses (1-2 vs 3-4) between the Confidence and Follow conditions to eliminate low-level visuomotor confounds (Figure 4D).

In independent across-domain classifications, we used the run-wise beta images reflecting JR- and CLR-activity as pattern vectors in a linear support vector classification model (as implemented in LIBSVM). We assigned each vector from each domain a label corresponding to the classes Confidence (1) and Follow (–1) in the JR-analysis and Low Confidence (–1) and High Confidence (1) in the CLR-analysis. We trained an SVM with the vectors from one domain (3 per class, 6 in total) and tested the decoder on the 6 vectors from the other domain (and vice versa) (Figure 4A; left), obtaining a mean average classification accuracy value for each of these two-way cross-classifications.

For within-domain classifications, we ran independent leave-one-run-out crossvalidations for each domain on JR-activity patterns (Confidence vs Follow) and CLR-activity patterns (Low vs High confidence). Pattern vectors from two of the three runs in each domain were used to train an SVM to predict the same classes in the vectors from the left-out run. We compared the true labels of the left-out run with the labels predicted by the model and iterated this process for the other two runs to calculate a mean cross-validated accuracy independently for each domain (Figure 4A; right).

We also tested the ability of confidence-related activity patterns to predict objective performance in the absence of confidence reports. We used a GLM that modeled Low vs High Confidence trials with a regressor that focused on the ‘rating period’, and Incorrect vs Correct Follow trials with a regressor that focused on the ‘decision period’ (i.e. from stimulus onset to subjects’ type I response). We performed a cross-classification analysis in which a decoder trained on Confidence trials (Low vs High Confidence) was tested on pattern vectors from Follow trials (Incorrect vs Correct), and vice versa (collapsed across domain). This confidence-objective performance generalization score was compared to a leave-one-run-out crossvalidation analysis decoding Low vs High confidence on Confidence trials only (collapsed across domain). Together, these scores characterize whether a particular set of patterns are specific to confidence, or also generalize to predict objective performance (Cortese et al., 2016) (Figure 5).

### Individual differences

Metacognitive efficiency scores (log meta-*d*’/*d*’) for each subject were estimated independently for the perceptual and memory tasks, together with a single score collapsed across domains. These scores were inserted as covariates in second-level analyses of within-perception, within-memory, and across-domain classifications of confidence level-related activity, respectively, to assess the parametric relationship between metacognitive efficiency and decoding success.

## Results

We analyzed the data of 24 subjects who underwent hemodynamic neuroimaging while performing two-alternative forced-choice discrimination tasks in perceptual (per) and memory (mem) domains (Figure 1A). In the perceptual task, subjects were asked to indicate the brighter of two stimuli (words or abstract shapes). In the memory task, subjects were asked to memorize exemplars of the same stimulus types, and then select the previously-learned stimulus from two stimuli presented on each trial. In half of the trials (“Confidence” condition) subjects performed a metacognitive evaluation after the discrimination task by rating their confidence in the correctness of their response by selecting a number on a 1-to-4 scale (1=not confident; 4=very confident). In order to differentiate metacognitive judgment-related activity from visuomotor activity engaged by use of the confidence scale, in the other half of trials (“Follow” condition) subjects were asked to respond according to a highlighted number without evaluating confidence in their response. To avoid stimulus-type confounds, two different types of stimulus—words and abstract shapes—were used as stimuli in both tasks.

### Behavior

We first compared task performance, measured by percentage of correct responses, across condition, task, and stimulus type. A 2×2×2 repeated measures ANOVA (confidence/follow × perception/memory × shapes/words) showed that performance was well-matched across conditions (Confidence vs Follow) (*F*_1,23_=3.036, *P*=0.095). None of the four paired *t*-tests (domain×stimulus) comparing performance between the Confidence and Follow conditions returned a significant difference (*P*>0.05). In the remainder of the behavioral analyses we focus on the Confidence condition. Matching performance across stimulus type was more challenging because subjects’ memory for words was expected to be considerably higher than that for abstract shapes trials based on pilot data (see Methods for details). Instead, we aimed to match subjects’ performance independently for each stimulus type across task domains by titrating the difficulty of the perceptual task to approximate the performance expected for the corresponding stimulus type in the memory task (shapes: per M=73%, mem M=67%; words: per M=81%, mem M=89%; Figure 1B). Critically, this ensured performance was matched across task domains when averaging stimulus types across participants (per: M=77%, mem: M=78%; paired *t*-test *T*_23_=0.38, *P*=0.70; Figure 1C). A 2×2 repeated measures ANOVA of performance in the Confidence condition (perception/memory × shapes/words) confirmed there was no main effect of domain (*F*_1,23_= 0.15, *P*=0.702). However, we observed a main effect of stimulus type due to greater overall performance on words (*F*_1,23_=75.69, *P*= 9.87× 10^−9^) and a domain × stimulus interaction, due to a greater difference in performance between shapes and words in the memory compared to the perception task (*F*_1,23_=16.74, *P*=0.00045).

Subjects were faster providing type I responses in perceptual trials (M=636ms) than in memory trials (M=1222ms). There was also a small difference in reaction times between shape (M=967ms) and word (M=892ms) trials. A 2×2 repeated measures ANOVA confirmed a main effect of domain (*F*_1,23_=367, *P*=1.23×10^−15^) driven by slower reaction types in the memory task. There was also a main effect of stimulus type on response time (*F*_1,23_=8.95, *P*=0.006), as well as a significant domain × stimulus interaction (*F*_1,23_=5.82,*P*=0.024).

As expected, subjects gave higher confidence ratings after correct decisions than after incorrect decisions (Figure 2A), and mean confidence ratings were similar across task domains (per M=2.62, mem M=2.47; paired *t*-test *T*_23_=1.26, *P*=0.22). Reaction times for confidence ratings were not different between domains (per M=518ms, mem M=516ms; paired *t*-test *T*_23_=0.16, *P*=0.87). We next estimated log (meta-*d*’/*d*’), a metacognitive efficiency measure derived from signal detection theory that assays the degree to which confidence ratings distinguish between correct and incorrect trials (Maniscalco and Lau, 2012; Fleming and Lau, 2014; Fleming, 2017). We used hierarchical Bayesian estimation to incorporate subject-level uncertainty in group-level parameter estimates (Fleming, 2017). Metacognitive efficiency in the perceptual task was significantly lower than in the memory task (*P_θ>0_ ~* 1; see Figure 2B & Methods for details), consistent with previous findings (Fleming et al., 2014). Metacognitive efficiency above optimality (meta-*d*’=*d*’) in memory trials suggests subjects had better metacognition than expected given their task performance, while suboptimal metacognitive efficiency in perceptual trials suggests subjects had worse metacognition than expected given their task performance (assuming an ideal observer in both cases). We did not find a correlation between subjects’ individual metacognitive efficiency scores in the perceptual and memory domains (r_22_=-0.076; *P*=0.72; Figure 2C). We also evaluated the correlation coefficient within a hierarchical model of meta-*d*’, which takes into account uncertainty in subject-level model fits (Fleming, 2017). The 95% confidence interval on the posterior correlation coefficient overlapped zero in this analysis (ρ=0.205; HDI=[0.826, −0.358]), also indicating a dissociation between domains.

**Figure 2.**
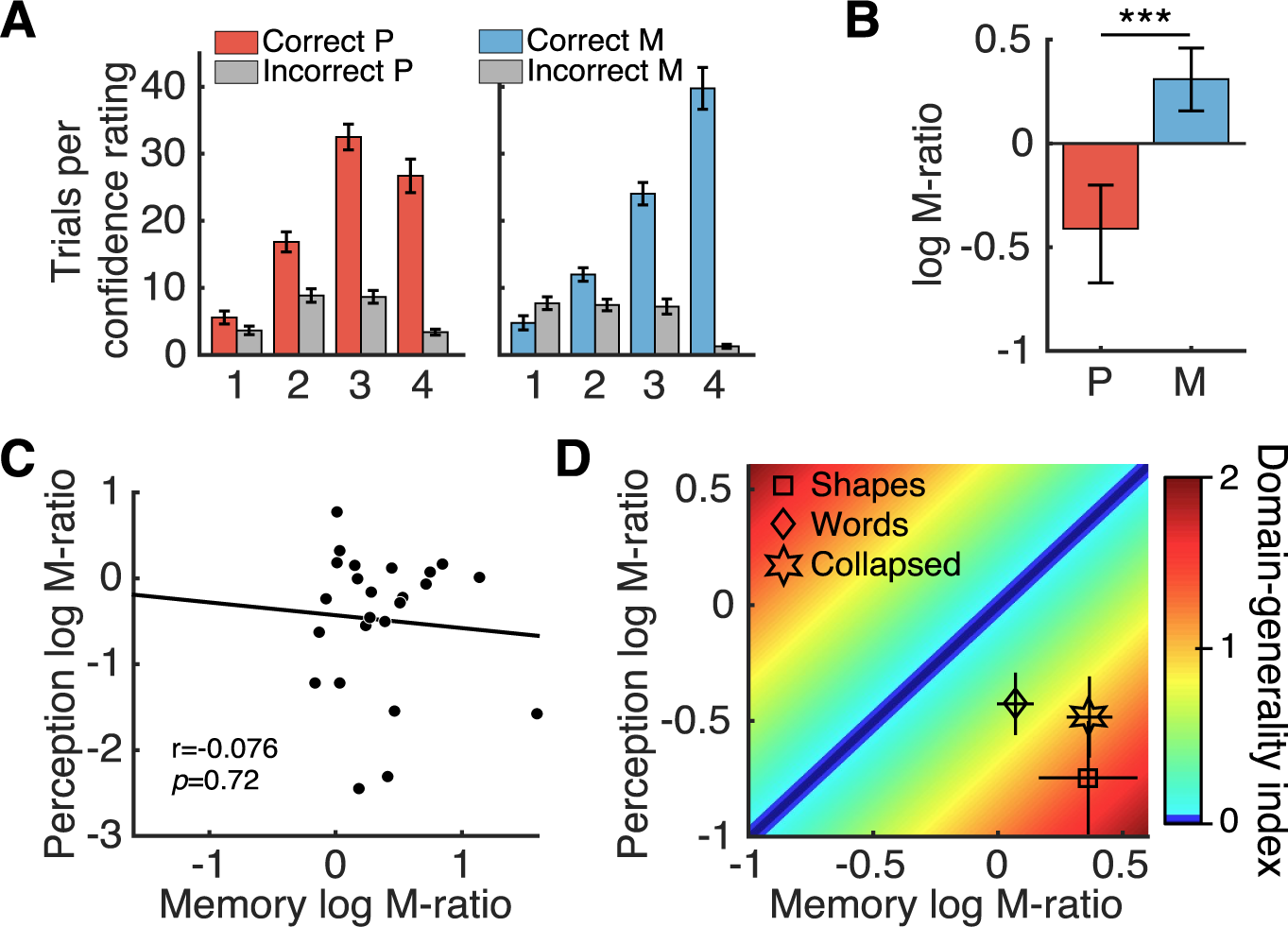
Metacognitive measures. **(A)** Mean number of correct and incorrect trials per confidence rating. **(B)** Metacognitive efficiency measured by log(meta-*d*’/*d*’). Zero indicates that metacognitive sensitivity (meta-*d*’) is equal to task sensitivity *d*’ (i.e. the *d*’ that would have been predicted to give rise to the observed confidence rating data assuming a signal detection theoretic ideal observer). Group-level hierarchical Bayesian estimates differed significantly between domains. Error bars indicate 95% high-density interval (HDI) from posterior samples. **(C)** Metacognitive efficiency scores obtained from single-subject Bayesian model fits were not correlated across perceptual and memory domains. **(D)** Domain-general index (DGI) for each subject that quantifies the similarity between their metacognitive efficiency scores in each domain (see main text for details). Greater DGI scores indicate less metacognitive consistency across domains. Mean log(meta-*d*’/*d*’) values for each stimulus type in both domains are shown for reference. Bars in (A) and (D) indicate s.e.m. *** *P*_*θ*>0_ ~ 1. P=perception M=memory.

We next estimated metacognitive efficiency separately for each stimulus type (Figure 2D). A 2×2 repeated measures ANOVA (perception/memory × shapes/words) indicated metacognitive efficiency was greater for memory than perception (*F*_1,23_=22.44, *P*=8.97×^10-5^). Importantly, there was no stimulus main effect (*F*_1,23_=0.015, *P*=0.902) and there was no interaction between domain and stimulus type (*F*_1,23_=2.835, *P*=0.106). To further assess a potential covariation between metacognitive abilities in each domain, we calculated for each subject a domain-generality index (DGI) that quantifies the similarity between scores in each domain for each participant (Fleming et al., 2014):

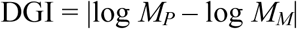

where *M_P_* = perceptual meta-*d*’/*d*’ and *M_m_* = memory meta-*d*’/*d*’. Lower DGI scores indicate more similar metacognitive efficiencies between domains (DGI=0 indicates identical scores). Mean DGI for shapes (1.42), words (0.66) and collapsed by stimulus type (0.95) were higher than zero (Figure 2D). Metacognition for words was behaviorally more stable across domains as the DGI was smaller than for shapes (paired *t*-test: *T*_23_=2.86; *P*=0.009). Together, these results suggest domain-specific constraints on metacognitive ability.

### fMRI analyses

We next turned to our fMRI data to assess the overlap between neural substrates engaged when metacognitive judgments are made during perceptual and memory tasks. A full understanding of the neural substrates of metacognition requires an independent examination of the process of engaging in a metacognitive task and the level of confidence expressed by the subject (Chua et al., 2014). To this end, in both univariate and multivariate analyses, we focused on two distinct features of metacognition-related activity. First, we assessed brain regions engaged in judgment-related (JR) activity (i.e. the difference between Confidence trials requiring a metacognitive judgment and the Follow condition). Second, we assessed brain regions engaged in confidence level-related (CLR) activity. In univariate CLR-analysis, we focused on the parametric relationship between confidence ratings (1 through 4) and neural activity. In multivariate CLR-analysis, we collapsed ratings 1 and 2 into a low-confidence category and ratings 3 and 4 into a high-confidence category to allow binary classification of activity patterns.

#### Univariate results

##### Judgment-related activity

In standard univariate analyses, we found elevated activity in dACC/pre-SMA, bilateral insulae and superior and middle frontal gyri when contrasting the Confidence against the Follow condition (collapsed by domain), consistent with previous findings (Fleming et al., 2012b) (Figure 3A). There were no significant clusters of activity in the reverse contrast (Follow>Confidence). Splitting the data by domain (see Table 1), an interaction contrast [Memory Confidence > Memory Follow] > [Perception Confidence > Perception Follow] revealed significant clusters of activity in middle cingulate gyrus, left insula, precuneus, left hippocampus and cerebellum (Figure 3B, blue). No significant clusters of activity were found in the reverse interaction contrast. In a conjunction analysis, elevated activity for the Confidence > Follow condition was observed across both perception and memory trials in anterior cingulate and right insula (Figure 3B, green).

**Figure 3.**
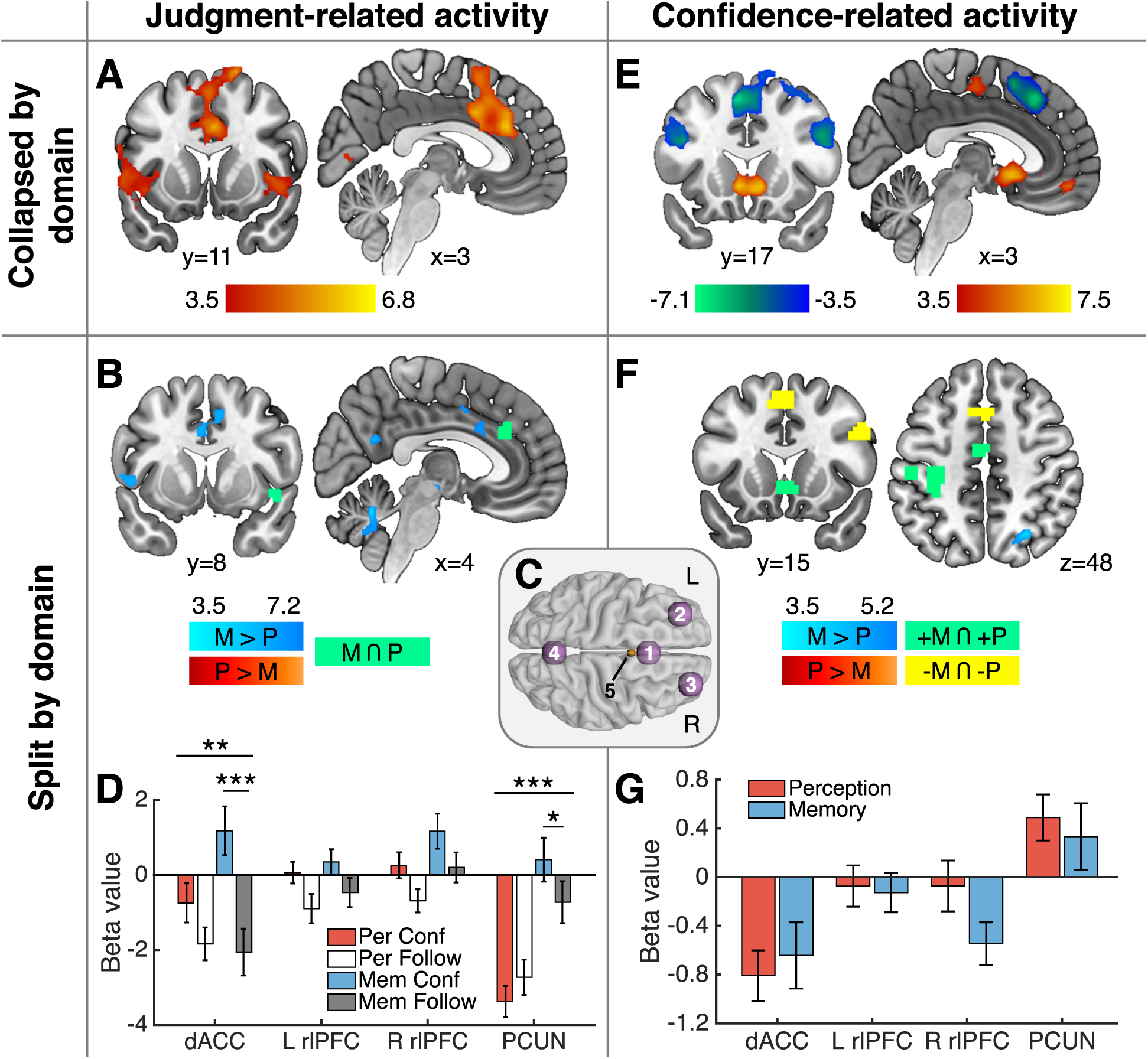
fMRI univariate analysis results.*Judgment-related activity*. **(A)** Whole-brain analysis of significant activation in the Confidence>Follow contrast (collapsed by domain); there were no significant clusters in the Follow>Confidence contrast. **(B)** [Memory Confidence > Memory Follow] > [Perception Confidence > Perception Follow] interaction contrast (blue). There were no significant clusters in the reverse contrast. The conjunction of Memory Confidence > Memory Follow and Perception Confidence > Perception Follow contrasts is indicated in green. **(C)** Spherical binary masks of four a priori regions of interest [1=dACC, 2=L rlPFC, 3=R rlPFC, 4=PCUN] and an ROI in the ventricles [5] used as a control region in multivariate analyses (Figure 4). **(D)** Estimated mean beta values for judgment-related activity by domain in the main four ROIs displayed in (C). *Confidence level-related activity*: **(E)** Whole-brain analysis of activity parametrically modulated by level of confidence (collapsed by domain). Hot colors indicate a positive correlation with confidence and cool colors a negative correlation. **(F)** Memory > Perception contrast (blue) testing for differences between the parametric effect of confidence by domain; there were no significant clusters in the Perception > Memory contrast. A conjunction analysis revealed shared activity that was positively (green) and negatively (yellow) correlated with confidence levels in both domains. **(G)** Estimated mean beta values of confidence level-related activity in the main four ROIs displayed in (C). All displayed whole-brain activations are significant at a cluster-defining threshold *P*<0.001, corrected for multiple comparisons *P_FWE_*<0.05; except for conjunction analyses in which we computed the intersection of two independent maps thresholded at *P*<0.001, uncorrected. Images displayed at *P*<0.001. Graded color bars reflect T-statistics. Error bars indicate s.e.m. *** *P*<0.001 ** *P*<0.01 \**P*<0.05. L=left; R=right; P=perception; M=memory.

**Table 1.**
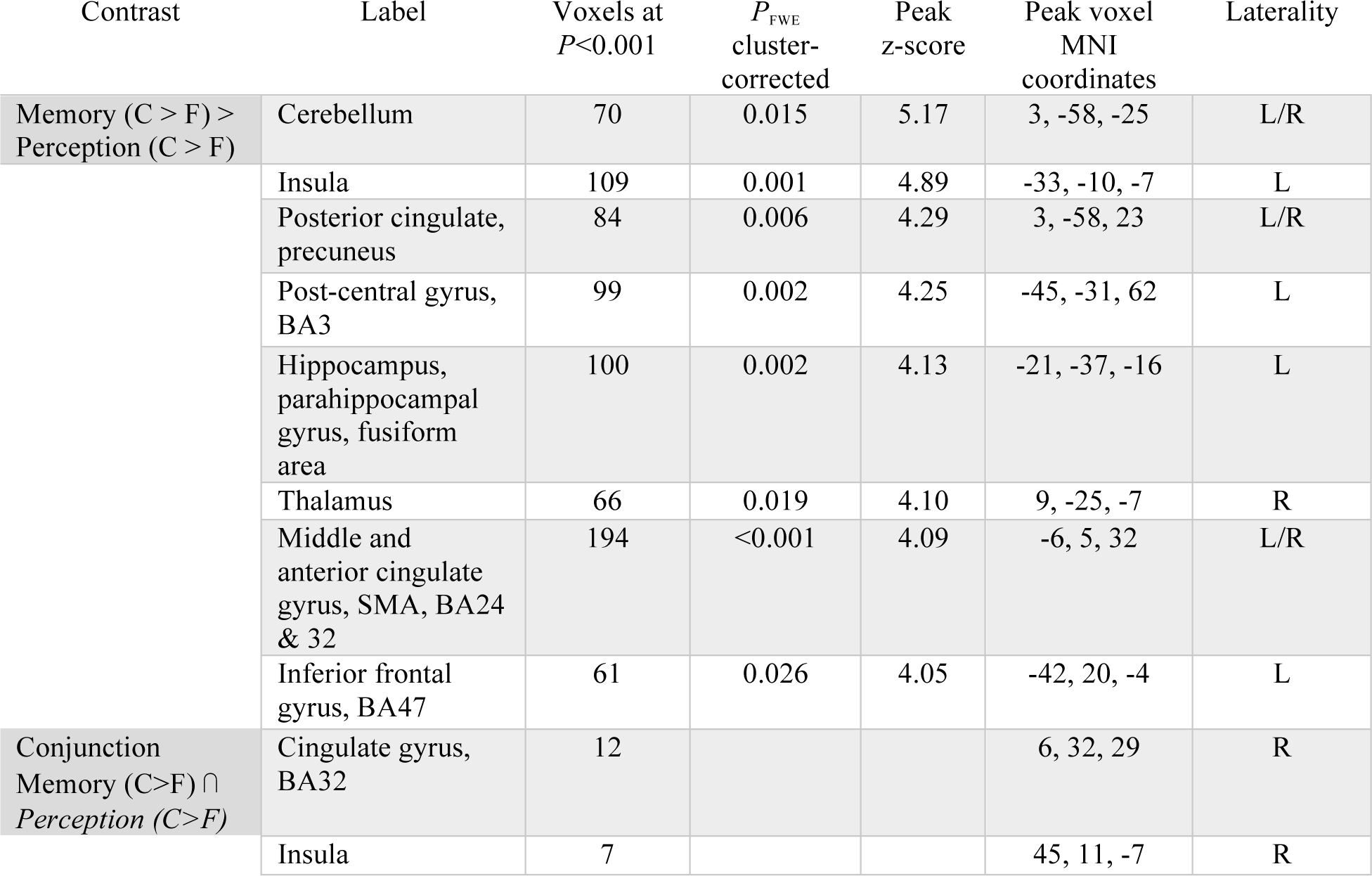
Univariate fMRI analysis - judgment-related activity interacted with domain.Significant activations at cluster-defining threshold *P*<0.001, corrected for multiple comparisons at *P*_FWE_<0.05. Conjunction of significant activations at cluster-defining threshold *P*<0.001, uncorrected, of Memory (C>F) and Perception (C>F) contrasts. See Figure 3B. C=confidence, F=follow

| Contrast | Label | Voxels at $P < 0.001$ | $P_{FWE}$ cluster-corrected | Peak z-score | Peak voxel MNI coordinates | Laterality |
| --- | --- | --- | --- | --- | --- | --- |
| Memory (C > F) > Perception (C > F) | Cerebellum | 70 | 0.015 | 5.17 | 3, -58, -25 | L/R |
|  | Insula | 109 | 0.001 | 4.89 | -33, -10, -7 | L |
|  | Posterior cingulate, precuneus | 84 | 0.006 | 4.29 | 3, -58, 23 | L/R |
|  | Post-central gyrus, BA3 | 99 | 0.002 | 4.25 | -45, -31, 62 | L |
|  | Hippocampus, parahippocampal gyrus, fusiform area | 100 | 0.002 | 4.13 | -21, -37, -16 | L |
|  | Thalamus | 66 | 0.019 | 4.10 | 9, -25, -7 | R |
|  | Middle and anterior cingulate gyrus, SMA, BA24 & 32 | 194 | <0.001 | 4.09 | -6, 5, 32 | L/R |
|  | Inferior frontal gyrus, BA47 | 61 | 0.026 | 4.05 | -42, 20, -4 | L |
| Conjunction Memory (C>F) ∩ Perception (C>F) | Cingulate gyrus, BA32 | 12 | | | 6, 32, 29 | R |
|  | Insula | 7 |  |  | 45, 11, -7 | R |

To further quantify these effects for each task domain, we focused on four a priori regions of interest (ROI) in (dACC/pre-SMA), bilateral rostrolateral prefrontal cortex (rlPFC) and precuneus (PCUN) (see Figure 3C & Methods), which previous studies have found to be recruited by perceptual and memory metacognition (Fleck et al., 2006; Fleming et al., 2010; 2012b; Baird et al., 2013; McCurdy et al., 2013). In a series of repeated measures 2×2 ANOVAS (condition × task) we found a main effect of greater activity on Confidence compared to Follow trials in all ROIs except precuneus, where instead we observed a main effect of task, with increased activity on memory trials (Figure 3D; condition main effect: dACC/pre-SMA, F_1,23_=19.34, *P*=0.0002; left rlPFC, F_1,23_3=6.62, *P*=0.017; right rlPFC, F_1,23_3=9.28, *P*=0.006; PCUN: F_1,23_=0.40, *P*=0.532; task main effect: dACC/pre-SMA, F_1,23_3=2.33, *P*=0.14; left rlPFC, F_1,23_=0.95, *P*=0.34; right rlPFC, F_1,23_3=4.94, *P*=0.036; PCUN: F_1,23_3=36.78, *P*=3.47×10^−6^). We found that the difference between Confidence and Follow trials was greater in memory than in perception trials in dACC/pre-SMA, recapitulating the whole-brain results (condition × task interaction; F_1,23_=12.16, *P*=0.0019; paired *t*-test, mem: T_23_=5.47, P=0.0001, per: T_23_=1.92, *P*= 0.067). A similar interaction pattern was observed in precuneus (condition × task interaction: F_1,23_=15.86, *P*=0.0006; paired *t*-test, mem: T_23_=2.43, *P*=0.023, per: T_23_=-1.54, *P*=0.136). There were no interactions in frontal regions (left rlPFC, F_1,23_=0.07, *P*=0.795; right rlPFC, F_1,23_=0.002, *P*=0.968). These results are compatible with previous findings indicating a distinctive contribution of precuneus to memory metacognition (Baird et al., 2013; McCurdy et al., 2013).

##### Confidence level-related activity

We next sought to investigate the parametric relationship between confidence level and neural activity. Collapsing across domains, we found activity in the left pre- and post-central gyri, the posterior midline, ventral striatum and ventromedial prefrontal cortex (vmPFC) correlated positively with confidence ratings (Figure 3E, hot colors). We also replicated negative correlations between confidence and activation in dACC/pre-SMA, parietal cortex, and bilateral PFC that have been reported in several previous studies (Fleck et al., 2006; Fleming et al., 2012b; Baird et al., 2013; Hebart et al., 2014) (Figure 3E, cool colors). When testing for differences between these parametric regressors by domain (see Table 2), a Memory > Perception contrast revealed a significant cluster of activity in right parietal cortex (Figure 3F, blue), while there was no significant activity in a Perception > Memory contrast. Shared positive correlations between confidence and activity in perception and memory trials were found in ventral striatum and in left pre- and post-central gyri, the latter consistent with use of the right hand to provide confidence ratings (conjunction analysis; Figure 3F, green). Shared negative correlations with confidence were found in regions of right dorsolateral PFC and medial prefrontal cortex, overlapping with pre-SMA (Figure 3F, yellow).

**Table 2.** Univariate fMRI analysis - confidence level-related activity interacted with domain. Significant activations at cluster-defining threshold *P*<0.001, corrected for multiple comparisons at *P_FWE_*<0.05. Conjunction of significant activations at cluster-defining threshold *P*<0.001, uncorrected, of Memory (M) and Perception (P) contrasts of positive and negative correlations with confidence level. See Figure 3F.

| Contrast | Label | Voxels at $P < 0.001$ | $P_{FWE}$ cluster-corrected | Peak z-score | Peak voxel MNI coordinates | Laterality |
| --- | --- | --- | --- | --- | --- | --- |
| Memory > Perception | Precuneus, BA7 | 93 | 0.003 | 4.21 | 33, -70, 20 | R |
| Conjunction (+M ∩ +P) | Precentral & postcentral gyri, BA6, 4, 3 | 167 | | | -30, -25, 44 | L |
|  | Post-central gyrus | 27 |  |  | -27, -46, 59 | L |
|  | SMA, BA6 | 21 |  |  | -3, -10, 50 | L/R |
|  | Ventral striatum | 16 |  |  | 3, 11, 7 | L/R |
|  | Cerebellum | 7 |  |  | 9, -58, -13 | R |
|  | Post-central gyrus | 5 |  |  | -48, -22, 44 | R |
| Conjunction (-M ∩ -P) | Middle frontal gyrus | 29 | | | 45, 26, 20 | R |
|  | Pre-SMA, BA8 | 19 |  |  | 0, 14, 50 | L/R |

**Table 3.**
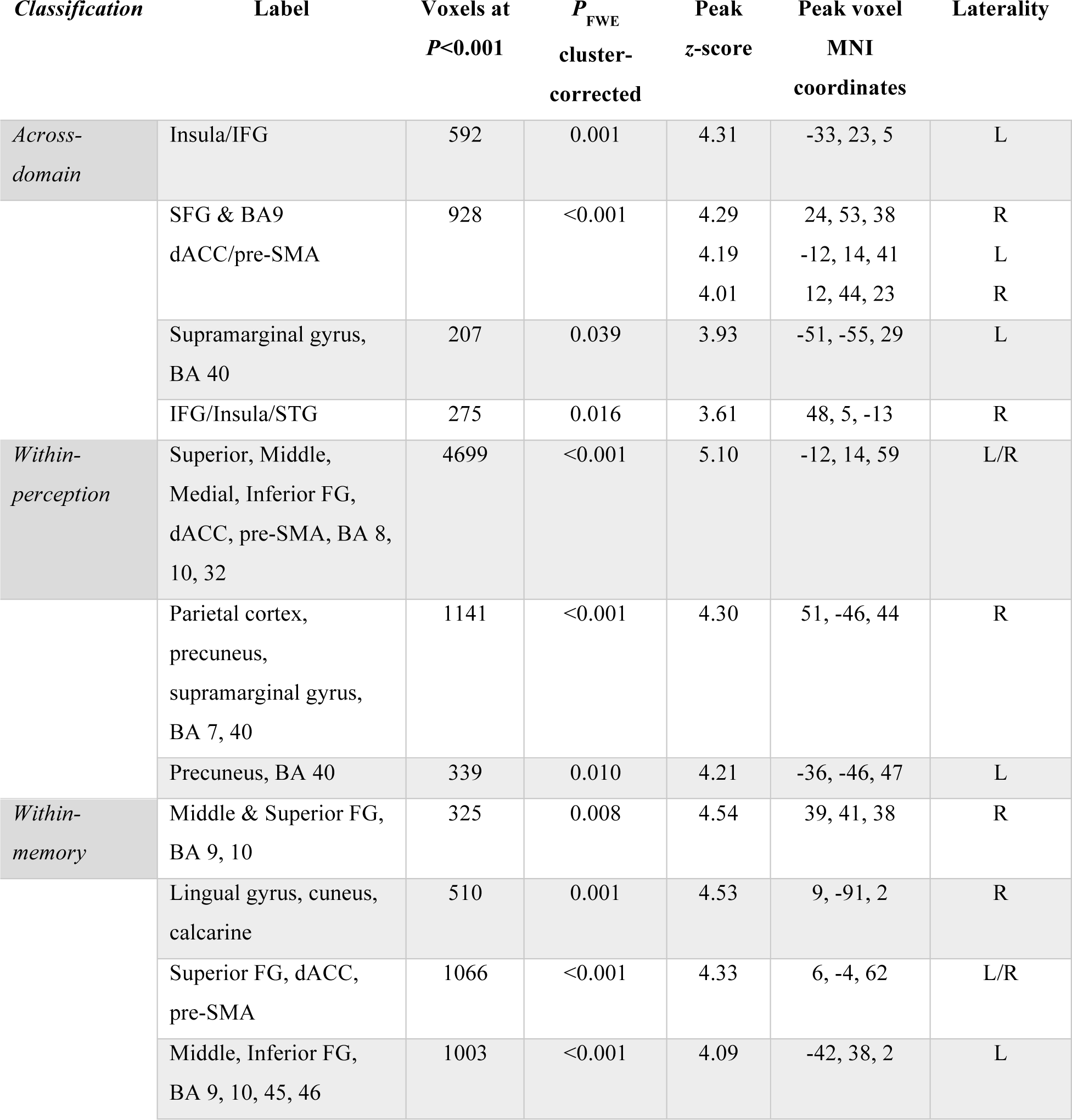
Judgment-related activity obtained from whole-brain searchlight classification analyses. Significant activations at cluster-defining threshold *P*<0.001, corrected for multiple comparisons at *P*_FWE_<0.05. See Figure 4C.

**Table 4.** Confidence level-related activity obtained from whole-brain searchlight classification analyses. Accuracy maps were masked to exclude visuomotor-related activity (see Figure 4D). Significant activations at cluster-defining threshold *P*<0.001, corrected for multiple comparisons at *P*_FWE_<0.05. See Figure 4F.

| *Classification* | *Label* | *Voxels at*<br>*$P < 0.001$* | *$P_{FWE}$*<br>*cluster-*<br>*corrected* | *Peak*<br>*z-score* | *Peak voxel*<br>*MNI*<br>*coordinates* | *Laterality* |
| --- | --- | --- | --- | --- | --- | --- |
| *Across-domain* | Inferior FG, ventral striatum, ACC, BA 11, 32 | 2434 | <0.001 | 5.20 | -21, 44, 2 | L/R |
|  | Superior and middle temporal gyri | 176 | <0.001 | 4.99 | -57, -52, 20 | L |
|  | Middle temporal gyrus, anterior cerebellum | 1193 | <0.001 | 4.92 | 3, -46, -34 | R |
|  | Middle cingulate gyrus | 46 | 0.017 | 4.85 | -6, -10, 47 | L/R |
|  | Superior temporal gyrus | 142 | <0.001 | 4.68 | 66, -7, 5 | R |
|  | Precuneus, BA 7, 19 | 543 | <0.001 | 4.68 | -12, -61, 65 | L/R |
|  | Superior temporal gyrus | 110 | <0.001 | 4.68 | -42, 26, -28 | L |
|  | Posterior cerebellum | 96 | <0.001 | 4.54 | -9, -85, -22 | L |
|  | SMA, BA 6 | 84 | 0.001 | 4.53 | -9, -10, 62 | L/R |
|  | Superior FG, dACC, pre-SMA | 482 | <0.001 | 4.39 | -3, 11, 56 | L/R |
|  | Post-central gyrus | 186 | <0.001 | 4.33 | -57, -19, 23 | L |
|  | Middle FG | 143 | <0.001 | 4.28 | 36, 5, 26 | R |
|  | Fusiform and parahippocampal gyri | 63 | 0.003 | 4.27 | 36, -19, -34 | R |
|  | Middle cingulate cortex | 64 | 0.003 | 4.25 | 0, -31, 53 | L/R |
|  | Posterior cerebellum | 53 | 0.008 | 4.10 | -48, -58, -37 | L |
|  | Inferior FG | 38 | 0.039 | 3.84 | 27, 23, -25 | R |
| *Within-perception* | Superior FG, dACC/pre-SMA | 185 | 0.032 | 3.88 | -6, 29, 47 | L/R |
| *Within-memory* | Inferior and middle FG, insula | 503 | <0.001 | 4.35 | 57, 23, 14 | R |
|  | dACC, pre-SMA | 211 | 0.021 | 3.79 | -9, 2, 47 | L/R |

Complementing the ROI analysis of JR-activity, we performed an ROI analysis of CLR-activity that recapitulated the whole-brain results. We observed negative relationships between confidence and activity in dACC/pre-SMA and positive relationships in precuneus. Importantly, no significant differences in the parametric effect of confidence were found between domains in any of our a priori ROIs (Figure 3G; paired *t*-tests: dACC/pre-SMA, T_23_=-0.47, *P*=0.643; left rlPFC, T_23_=0.23, *P*=0.820; right rlPFC, T_23_=1.62, *P*=0.119; PCUN: T_23_=0.56, *P*=0.583). Together with the lack of marked domain-specific differences in confidence-related activity at the whole-brain level, these results are suggestive of an absence of domain-specificity in confidence-related activity. However, a lack of difference between univariate activation profiles is not necessarily conclusive. For instance, differences in confidence level may be encoded in fine-grained spatial patterns of activity even when the overall BOLD-activity is evenly distributed across confidence levels (Cortese et al., 2016). Similarly, while metacognition-related activity may show similar overall levels of activation across tasks, distributed activity patterns in frontal and parietal areas may carry distinct task-specific information (Hebart et al., 2014; Cole et al., 2016). We next turned to multivariate analysis methods, which are sensitive to differences in spatial activity patterns, to test this hypothesis.

#### Multivariate results

We performed a series of multi-voxel pattern analyses (MVPA) (Figure 4A) focused on both judgment-related activity patterns and confidence level-related activity patterns.

**Figure 4.**
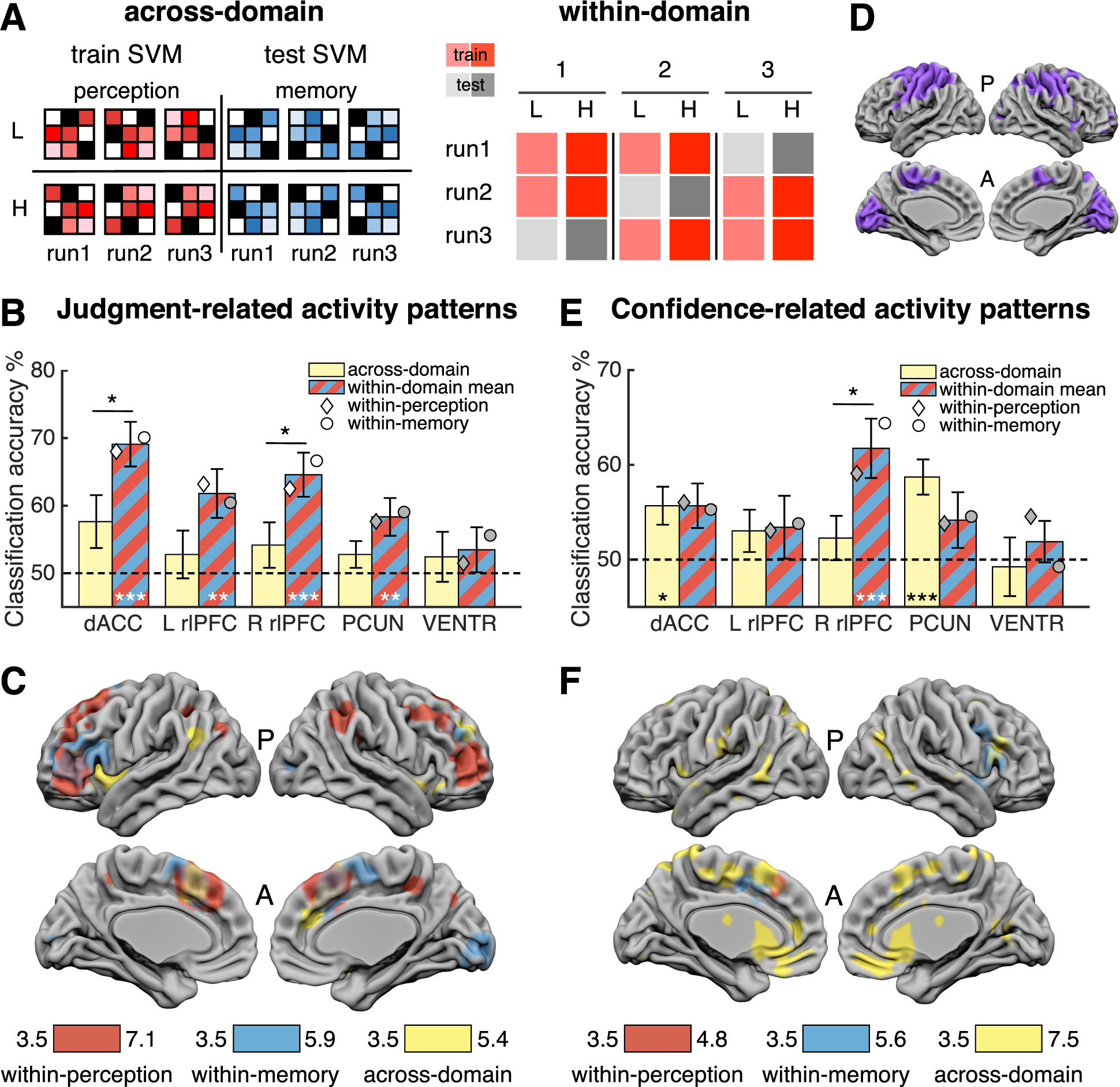
MVPA Results. *Classification designs*. **(A)** *Left*: Across-domain classification design. Pattern vectors (run-wise beta images) from one domain were used to train an SVM decoder on two classes and then tested in a cross-classification of the same two classes using vectors from the other domain (and vice versa). Classification of low (L) and high (H) confidence levels is illustrated here. *Right:* Within-domain classification design. Pattern vectors of two classes (e.g. low and high confidence) pertaining to one domain were used to train a decoder in a leave-one-run-out design that was then tested in the left-out pair. The process was iterated three times to test pairs from every run. An identical, independent cross-validation was performed on vectors from the other domain. *Judgment-Related Activity Patterns:* **(B)** ROI results for across-domain (yellow) and mean within-domain (red-blue stripe) classification accuracy of Confidence vs Follow trials. **(C)** Searchlight analysis for same classifications as in (B). **(D)** Low-level visuomotor mask used in (F) (see main text and Methods for details). *Confidence Level-Related Activity Patterns: (E)* Low versus high confidence classification accuracy results. **(F)** Searchlight analysis for the same classifications as in (E) exclusively masked for visuomotor-related activity patterns. Bars in (B) and (E) indicate means and error bars s.e.m. Dashed lines indicate chance classification (50%). Diamonds and circles indicate mean independent classification in perception and memory trials, respectively. White diamonds/circles indicate classification was significantly different from chance, Bonferroni corrected. All clusters in (C) and (F) are significant at cluster-defining threshold *P*<0.001, corrected for multiple comparisons at *P_FWE_*<0.05. Image displayed at *P*<0.001. Color bars indicate T-scores. P=perception; M=memory; A=anterior; P=posterior. *** *P*≤0.001 ** *P*≤0.01 * *P*<0.05; all one-sample *t*-tests are Bonferroni corrected.

##### ROI analysis of judgment-related activity patterns

If metacognitive judgments are based on domain-general processes (i.e. shared across perceptual and memory tasks), a decoder trained to classify JR-activity patterns in perceptual trials should accurately discriminate JR-activity patterns when tested on memory trials (and vice versa). Alternatively, domain-specific activity profiles would be indicated by significant within-domain classification of JR-activity patterns in the absence of across-domain transfer. To adjudicate between these hypotheses, we performed a support vector machine (SVM) decoding analysis using as input vectors the run-wise beta images pertaining to Confidence and Follow trials obtained from a GLM (12 input vectors in total). For within-domain classification we used standard leave-one-out independent crossvalidations for each domain and we tested for across-domain generalization using a cross-classification analysis (see Methods for details). Chance classification in both analyses was 50%.

Mean within-domain classification results were significantly above chance in all ROIs (one-sample *t*-tests Bonferroni corrected for multiple comparisons α=0.05/4=0.0125: dACC/pre-SMA *T*_23_=5.77, *P*=6.99×10^−6^; L rlPFC *T*_23_=3.27, *P*=0.003; R rlPFC *T*_23_=4.47, *P*=0.0002; PCUN *T*_23_=2.98, *P*=0.007; Figure 4B, red-blue stripe bars). In contrast, across-domain generalizations were not significantly different from chance in any ROI (one sample *t*-test Bonferroni corrected: dACC/pre-SMA, *T*_23_=1.95, *P*=0.06; L rlPFC, *T*_23_=0.79, *P*=0.44; R rlPFC, *T*_23_=1.24, *P*=0.23; PCUN *T*_23_=1.40, *P*=0.17; Figure 4B, yellow bars). As a control, classification accuracy in the ventricles (VENTR) was not different from chance (across: *T*_23_=0.66, *P*=0.52; within: *T*_23_=1.04, P=0.31). This suggests that the patterns of activity that distinguish metacognitive judgments from the visuomotor control condition in one domain are distinct from analogous patterns in the other domain. In particular, within-domain classification accuracy was significantly different from across-domain classification accuracy in (dACC/pre-SMA: *T*_23_=2.88, *P*=0.008) and right rlPFC (*T*_23_=2.24, *P*=0.035). These results are consistent with the hypothesis that metacognitive judgments recruit domain-specific patterns of cortical activity in prefrontal cortex.

##### Searchlight analysis ofjudgment-related activity patterns

We ran a similar decoding analysis using an exploratory whole-brain searchlight, obtaining a classification accuracy value per voxel (Hebart et al., 2015). Consistent with our ROI results, we observed significant within-domain classification in large swathes of bilateral PFC for both perception (red) and memory (blue) (Figure 4C). Within-perception classification was also successful in parietal regions—precuneus in particular—and within-memory activity patterns were classified accurately in occipital regions. We also identified clusters showing significant across-domain generalization (yellow) in dACC, pre-SMA, SFG (BA 9), supramarginal gyrus (BA 40), and bilateral IFG/insula, consistent with univariate results (Figure 3 A).

##### ROI analysis of confidence level-related activity patterns

We next asked whether confidence is encoded in a domain-general or domain-specific fashion by applying a similar approach to discriminate low versus high confidence trials. Note that in this case, ROI univariate analyses did not reveal any differences in confidence-related activity between domains (Figure 3G). We hypothesized that if confidence level is encoded by domain-general neural activity patterns, it should be possible to train a decoder to discriminate low (1-2) from high (3-4) confidence rating patterns in the perceptual task and then accurately classify confidence patterns on the memory task (and vice versa). In the absence of across-domain classification, significant within-domain classification is indicative of confidence level-related domain-specific activity patterns. ROI cross-classifications and cross-validations were performed in a similar fashion as above. We note that two subjects did not provide ratings for one of the classes in at least one run and were left out from the main analysis to avoid entering unbalanced training data into the classifier; however, including those subjects did not change the main result.

One-sample *t*-tests (Bonferroni-corrected) showed across-domain classification of confidence was significantly above chance in dACC/pre-SMA (*T*_21_=2.83, P=0.010) and precuneus (*T*_21_=4.69, *P*=0.0001), indicative of a generic confidence signal, but not in rlPFC (left: *T*_21_=1.36, *P*=0.19; right: *T*_21_=0.97, *P*=0.34) (Figure 4E, yellow bars). In contrast, mean within-domain classification accuracy was significantly above chance in right rlPFC (*T*_21_=3.75, *P*=0.001), but not in the other ROIs (dACC/pre-SMA: *T*_21_=2.42, *P*=0.025; left rlPFC: *T*_21_=1.03, *P*=0.32; PCUN: *T*_21_=1.42, *P*=0.17; Bonferroni-corrected; Figure 4E, red-blue stripe bars). The mean confidence classification accuracy in right rlPFC was 62% (per=59%, Mem=64%), notably above a recently estimated median 55% for decoding task-relevant information in frontal regions (Bhandari et al., 2017). Importantly, classification accuracy in this ROI also differed from the corresponding across-domain classification accuracy (paired *t*-test *T*_21_=2.37, *P*=0.028). Classification accuracy in the ventricles was not different from chance (across: *T*_21_=-0.24, *P*=0.81; within: *T*_21_=0.86, *P*=0.40). When subjects with unbalanced data were included, within-domain classification accuracy in right rlPFC remained at 62%, significantly above chance (*T*_23_= 4.22; *P*=0.0003) and significantly different from across-domain classification accuracy (paired *t*-test: *T*_23_= 2.54; *P*=0.018). Together, these results suggest the co-existence of two kinds of CLR-neural activity: dACC/pre-SMA and precuneus encode a generic confidence signal, whereas patterns of activity in right rlPFC were modulated by task, allowing within-but not across-domain classification of confidence level.

##### Searchlight analysis of confidence level-related activity patterns

We ran a similar decoding analysis of confidence level using an exploratory whole-brain searchlight, obtaining a classification accuracy value per voxel. Here we leveraged the Follow trials as a control for low-level visuomotor confounds by exclusively masking out activity patterns associated with usage of the confidence scale (Figure 4D). The remaining activity patterns can therefore be ascribed to confidence level-related signals that do not encode visual or motor features of the rating (Figure 4F). We found widespread across-domain classification of confidence (yellow) in a predominantly midline network including a large cluster encompassing dACC/pre-SMA, vmPFC, and ventral striatum. Domain-specific confidence level-related activity patterns were successfully decoded from right PFC (insula, IFG, BA 9, 46) in memory trials (blue) and were also independently decoded in both domains from dACC/pre-SMA.

##### Generalization of CLR-activity to objective performance

To further address the question of how confidence judgments may relate to activity patterns, we examined the relationship between objective task accuracy and confidence. Previous work suggests that the neural basis (and associated activation patterns) of confidence and performance may be partly distinct (Rounis et al., 2010; Cortese et al., 2016). Specifically, we tested the hypothesis that we could train a decoder using CLR-activity patterns to classify objective performance-related activity patterns (correct/incorrect) on Follow trials (and vice versa) in a cross-classification analysis (collapsed across domain). This analysis confirmed that activity patterns in dACC/pre-SMA (*T*_21_=2.38, *P*=0.027) and right rlPFC (*T*_21_=2.64, *P*=0.015) could predict objective accuracy levels in Follow trials above chance (Figure 5, light gray; uncorrected), but not in left rlPFC (*T*_21_=1.49, *P*=0.15) or precuneus (*T*_21_=−0.46, *P*=0.65). We then compared these decoding scores to a leave-one-run-out cross-validation decoding analysis of low versus high confidence on Confidence trials only (collapsed by domain; Figure 5, dark gray; uncorrected). Consistent with the analyses reported in Figure 4E (yellow), this decoder was unable to classify domain-general confidence patterns of activity in right rlPFC (*T*_21_=1.00; *P*=0.33), but was above chance in dACC/pre-SMA (*T*_21_=3.80, *P*=0.001), left rlPFC (*T*_21_=2.26, *P*=0.034) and precuneus (*T*_21_=2.56, *P*=0.018). Critically, in precuneus, confidence classification was significantly greater than confidence-performance generalization, which was at chance (paired *t*-test *T*_21_=2.16, *P*=0.043). This result indicates that confidence-related patterns in precuneus do not generalize to predict objective performance, consistent with a partly distinct coding of information relevant to task performance and confidence. In contrast, in dACC/pre-SMA, general confidence level and performance could be predicted from common patterns of activation.

**Figure 5.**
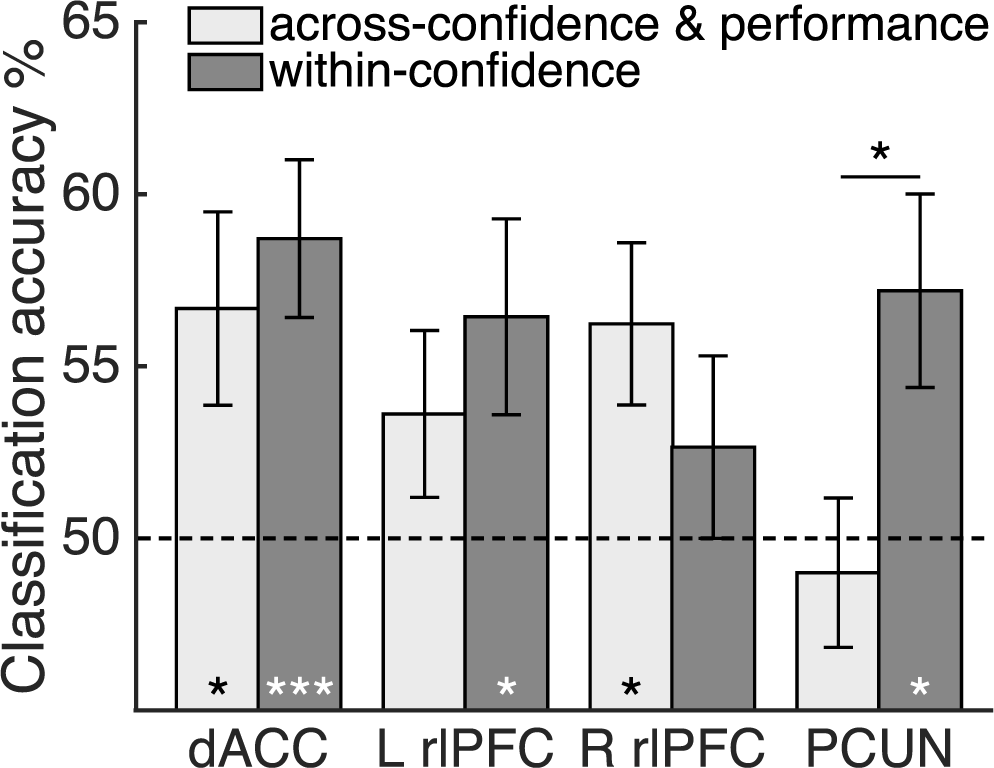
Generalization of confidence-related activity to objective accuracy. Light gray bars denote mean cross-classification accuracy results obtained from training a decoder on CLR-activity and testing it on objective accuracy (correct / incorrect) activity patterns in the Follow condition (and vice versa). Dark gray bars denote decoding accuracy for a leave-one-out crossvalidation of low and high confidence on Confidence trials only (collapsed by domain). Bars indicate group means and bars s.e.m. Dotted line indicates chance level. * *P*<0.05 *** *P*<0.001

##### Metacognitive efficiency and confidence level-related activity classification

Finally, we reasoned that if confidence-related patterns of activation are contributing to metacognitive judgments, they may also track individual differences in metacognitive efficiency. To test for such a relation, we asked whether individual metacognitive efficiency scores collapsed across-domains and independently in each domain predicted searchlight classification accuracy of confidence level. We did not find any significant clusters after whole-brain correction for multiple comparisons in domain-general, within-perception, or within-memory analyses. However, memory metacognitive efficiency predicted memory confidence classification accuracy in a cluster in right precuneus and left precentral gyrus (*P*<0.001, uncorrected), whereas perceptual metacognitive efficiency predicted perceptual confidence classification accuracy in left middle frontal gyrus, right vmPFC, bilateral temporal gyri, and cerebellum (*P*<0.001, uncorrected). Previous studies (Fleming et al., 2010; McCurdy et al., 2013; Fleming et al., 2014) have reported similar relations between perceptual and memory metacognitive efficiency and individual differences in the structure of prefrontal and parietal cortex, respectively. While we do not interpret these findings further in the current manuscript, for completeness, second-level unthresholded statistical images of these analyses were uploaded to Neurovault in order to inform future work (Gorgolewski et al., 2015) (https://neurovault.org/collections/3232/).

## Discussion

When performing a cognitive task, confidence estimates allow for comparisons of performance across a range of different scenarios (de Gardelle and Mamassian, 2014). Such estimates must also carry information about the task context if they are to be used in decision-making. Here we investigated the domain-generality and domain-specificity of representations that support metacognition of perception and memory.

Unlike previous studies (McCurdy et al., 2013), subjects’ performance was matched between domains for two different types of stimulus, thereby eliminating potential performance and stimulus confounds. Subjects’ confidence ratings were also matched between domains and followed expected patterns of higher ratings after correct decisions than after incorrect decisions. Metacognitive efficiency scores between tasks were not correlated, and metacognitive efficiency scores in the memory task were superior to those in the perceptual task. Using univariate and multivariate analyses, we showed the existence of both domain-specific and domain-general metacognition-related activity during perceptual and memory tasks. We report four main findings, and discuss each of these in turn.

First, we obtained convergent evidence from both univariate and multivariate analyses that a cingulo-opercular network centered on dACC/pre-SMA encodes a generic signal predictive of confidence level and objective accuracy across memory and perceptual tasks. Previous studies of metacognition have implicated the cingulo-opercular network in tracking confidence level (Fleck et al., 2006; Fleming et al., 2012b; Hebart et al., 2014; Hilgenstock et al., 2014). However, we go beyond these previous studies to provide evidence that these signals generalize to predict confidence across two distinct cognitive domains. This finding is consistent with posterior medial frontal cortex as a nexus for monitoring the fidelity of generic sensorimotor mappings, building on previous findings that error-related event-related potentials originating from this region are sensitive to variation in subjective certainty (Scheffers and Coles, 2000; Boldt and Yeung, 2015). The activity in dACC/pre-SMA was also consistently elevated by the requirement for a metacognitive judgment (Fleming et al., 2012b). However, the results regarding the generalizability of the pattern of these increases across tasks were inconclusive. Whole-brain searchlight analysis revealed successful cross-classification of these activity patterns in dACC and insular regions, consistent with the results of the univariate analysis. These patterns of activity, however, did not generalize across tasks in a pre-defined region of interest centered in dACC/pre-SMA.

Notably, while both dACC/pre-SMA and precuneus showed significant domain-general decoding of confidence, in precuneus these patterns did not generalize to also predict changes in objective accuracy. While performance and subjective confidence may both depend on similar decision variables (Kiani and Shadlen, 2009; Fleming and Daw, 2017), behavioral dissociations between these quantities are also consistent with distinct internal states contributing to decisions and confidence ratings (Busey et al., 2010; Fleming and Daw, 2017). For instance, hierarchical models of confidence formation suggest a downstream network “reads out” decision-related information in a distinct neural population (Insabato et al., 2010). The observed lack of crossclassification in precuneus (Figure 5) is consistent with the recent observation of distinct neural patterns of activity pertaining to confidence and first-order performance revealed through multivoxel neurofeedback in frontal and parietal regions (Cortese et al., 2016).

Second, in lateral anterior frontal cortex we found activity patterns that tracked both the requirement for metacognitive judgments and level of confidence. Large swathes of lateral prefrontal cortex distinguished activity patterns pertaining to metacognitive judgments that were specific for each domain. Critically, however, confidence-related activity patterns were selective for domain in right rlPFC (Figure 4E): they differed according to whether the subject was engaged in rating confidence about perception or memory. Such signals may support the “tagging” of confidence with contextual information, thereby facilitating the use of confidence for behavioral control (Donoso et al., 2014; Purcell and Kiani, 2016). The identity of perceptual and memory tasks can be reliably decoded from activity in right PFC neural populations (Mendoza-Halliday and Martinez-Trujillo, 2017), consistent with the possibility that this contextual information is recruited during confidence rating. It is possible that anterior prefrontal regions combine generic confidence signals with domain-specific information to fine-tune decision-making and action selection in situations in which subjects need to regularly switch between tasks or strategies on the basis of their reliability (Donoso et al., 2014). An alternative hypothesis, also compatible with our data, is that PFC first estimates the confidence level specifically for the current task, which is then relayed to medial areas to recruit the appropriate resources for cognitive control in a task-independent manner. Processing dynamics may also unfold simultaneously in both areas. These possibilities echo a longstanding debate in the cognitive control literature on the relative primacy of medial and lateral PFC in the hierarchy of control (Kerns et al., 2004; Tang et al., 2016). Further inquiry and development of computational models of the hierarchical or parallel functional coupling of these networks in metacognition is necessary.

Third, we obtained convergent evidence that precuneus plays a specific role in metamemory judgments. In univariate fMRI analyses, we found the requirement for a metacognitive judgment recruited our pre-established region of interest centered on precuneus only on memory, but not perceptual, trials (Figure 3D). Individual metacognitive efficiency scores in memory trials predicted classification accuracy in a more dorsal precuneal region, while individual differences in metacognitive efficiency scores in perceptual trials predicted classification accuracy in ventromedial prefrontal cortex (albeit at uncorrected thresholds). These findings are consistent with the medial parietal cortex making a disproportional contribution to memory metacognition (Simons et al., 2010; Baird et al., 2013; McCurdy et al., 2013), and offer a potential explanation for a decrease in perceptual, but not memory, metacognitive efficiency seen in patients with frontal lesions (Fleming et al., 2014). However, we do not wish to conclude that precuneus involvement is specific to metamemory. We note that univariate negative correlations with confidence were found also on perceptual trials, and multivariate classification results in precuneus indicated the presence of both perceptual and memory-related signals. This dual involvement of the precuneus in perception and memory metacognition is consistent with previous studies which suggest a relationship between precuneus structure and visual perceptual metacognition (Fleming et al., 2010; McCurdy et al., 2013).

Fourth, we found in both univariate and multivariate whole-brain analyses that domain-general signals in the ventral striatum and vmPFC (including subgenual anterior cingulate cortex) were modulated by confidence level. These results are compatible with previous findings that have found activity in ventral striatum to be positively correlated with confidence (Daniel and Pollmann, 2012; Hebart et al., 2014; Guggenmos et al., 2016). Evidence of vmPFC encoding of confidence signals has been reported in connection to decision-making and value judgments (De Martino et al., 2013; Lebreton et al., 2015). Our experimental design, however, does not allow us to disentangle whether the signals found in these regions pertain uniquely to confidence or whether they are entangled with implicit value and reward signals (for instance, the expected value of being correct). Future experiments are needed to explicitly decouple reward from confidence to resolve this issue.

In our experimental design perception and memory blocks were interleaved across runs, which raises the question as to whether the domain-specific neural substrates we found would persist if subjects had to switch between tasks more often. Due to inter-task “leaks” in confidence (Rahnev et al., 2015), in which confidence in one task influences confidence ratings on the following task (or even in the following trial), there is a possibility that interleaving blocks of different tasks might favor the observation of more domain-general confidence-related patterns.

Our experimental design assumes that visual perception and memory are distinct domains. We acknowledge that distinguishing between cognitive domains or individuating perceptual modalities is not straightforward (Macpherson, 2011). For instance, different modalities (e.g. vision, audition, touch, etc.), different aspects within a single modality (e.g. motion and color within vision), or closely related modalities (e.g. visual perception vs visual short-term memory) could be part of a unified perceptual domain for metacognitive purposes. Recent findings suggest metacognitive efficiency in one perceptual modality predicts metacognitive efficiency in others and that they share electrophysiological markers (Faivre et al., 2017). Metacognitive efficiency is also correlated across vision and visual short-term memory, especially for features such as orientation (Samaha and Postle, 2017), and dACC and insula regions similar to those identified here have been found to show univariate confidence signals across both color and motion tasks in the visual domain (Heereman et al., 2015). However, it is an open question whether more fine-grained modality-specific patterns of metacognitive activity could be decoded using multivariate approaches. More research is needed on the neural architecture of metacognition in other cognitive domains, and whether this architecture changes in a graded or discrete fashion as a function of task or stimulus.

In summary, our results provide evidence for the co-existence of content-rich metacognitive representations in anterior prefrontal cortex with generic confidence-related signals in frontoparietal and cingulo-opercular regions. Such an architecture may be appropriate for “tagging” lower-level feelings of confidence with higher-order contextual information to allow effective behavioral control. Previous studies have tended to draw conclusions about either domain-specific or domain-general aspects of metacognition. Here we reconcile these perspectives by demonstrating that both domain-specific and domain-general signals co-exist in the human brain, thus laying the groundwork for a mechanistic understanding of reflective judgments of cognition.

## Acknowledgements

We thank Aurelio Cortese for his help with the MVPA analyses. This work was partially supported by the National Institute of Neurological Disorders and Stroke of the National Institutes of Health (Grant No. R01NS088628) to H.L. and S.M.F., by the Templeton Foundation (Grant No. 21569) to H.L., and by a Sir Henry Wellcome Fellowship to S.M.F. (Grant No. WT096185). The Wellcome Centre for Human Neuroimaging is supported by core funding from the Wellcome Trust (Grant No. 203147/Z/16/Z).

